# A novel mechanism for morphological change during bacterial sporulation based on programmed peptidoglycan degradation

**DOI:** 10.1101/2025.06.26.661752

**Authors:** Carlos A. Ramírez Carbó, Oihane Irazoki, Srutha Venkatesan, Lauren J. S. Chen, Haylie A. Morales, Assariel J. Garcia Avila, Hoi-Ling Cheung, Felipe Cava, Beiyan Nan

**Affiliations:** Department of Biology, Texas A&M University, College Station, Texas, USA; The Genetics and Genomics Interdisciplinary Program, Texas A&M University, College Station, TX 77843, USA; Department of Molecular Biology and Laboratory for Molecular Infection Medicine Sweden, Umeå Centre for Microbial Research, SciLifeLab, Umeå University, Umeå, Sweden

## Abstract

Many bacteria form spores to endure unfavorable conditions. While *Firmicutes* generate endospores through cell division, sporulation in non-Firmicutes remains less understood. The Gram-negative bacterium *Myxococcus xanthus* undergoes sporulation through two distinct mechanisms: rapid sporulation triggered by chemical induction and slow sporulation driven by starvation, both occurring independently of cell division. Instead, these processes depend on the complete degradation of the peptidoglycan (PG) cell wall by two lytic transglycosylases (LTGs), LtgA and LtgB. Remarkably, LtgB programs the pace of PG degradation by LtgA during rapid sporulation, ensuring a controlled process that prevents abrupt PG breakdown and the formation of non-resistant pseudospores. In addition to regulation between LTGs, PG degradation is also influenced by its synthesis; cells exhibiting increased muropeptide production often circumvent sporulation. These findings not only reveal novel mechanisms of bacterial sporulation but also shed light on the regulatory network governing PG dynamics.

## Introduction

Spores are metabolically dormant cells that can survive unfavorable conditions, including extremes of temperature, desiccation, and ionizing radiation (Huang & Hull, 2017, Hutchison *et al*., 2014). Sporulation, the process of spore development, is a strategy utilized by a wide variety of organisms, from bacteria and protozoa to fungi and plants. In addition to protective spore coats, spore resilience relies on cytosol dehydration and DNA compaction—key processes that decrease cell volume and alter cell shape (Higgins & Dworkin, 2012). Thus, morphological differentiation is a hallmark of sporulation. In bacterial spore formers, as peptidoglycan (PG) cell walls largely determine cell morphology, their sporulation requires profound PG remodeling.

PG is a rigid, mesh-like macromolecule that is composed of glycan strands of repeating units of N-acetyl glucosamine (GluNAc)-N-acetyl muramic acid (MurNAc) crosslinked by peptides (Egan *et al*., 2020). PG encloses the entire cytoplasmic membrane, and its rigidity provides mechanical support against osmotic stress. For this reason, PG is an essential structure for most bacteria and a major target for antibacterial treatments. Sporulation provides invaluable opportunities for understanding the dynamics of PG, which is controlled by multiple synthases and hydrolases. PG synthases include glycosyltransferases (GTases) that polymerize glycan strands and transpeptidases (TPases) that form peptide crosslinks (Egan *et al*., 2020). PG hydrolases, also known as autolysins, include glycosidases that break the glycan strands, and amidases and endo/carboxypeptidases that cleave the peptide crosslinks (Egan *et al*., 2020, Rohs & Bernhardt, 2021, van Heijenoort, 2011).

*Firmicutes*, including Gram-positive bacteria like *Bacilli* and *Clostridia*, produce endospores. Their morphological transition from rod-shaped vegetative cells to ovoid spores occurs through asymmetric cell division, resulting in the formation of a smaller forespore and a larger mother cell. Eventually, the forespore becomes an oval endospore after being engulfed by the mother cell (Higgins & Dworkin, 2012, McKenney & Eichenberger, 2012). A mature endospore contains two PG layers: the germ cell wall, derived from the forespore’s original PG, and a thickened PG cortex deposited by the mother cell (Popham & Bernhards, 2015). While a few Gram-negative spore formers also belong to the *Firmicutes* phylum, they share conserved sporulation genes with *Bacilli* and *Clostridia*, which suggests similar sporulation mechanisms (Tocheva *et al*., 2011, Yutin & Galperin, 2013).

Sporulation by non-firmicutes bacteria has been largely overlooked. Myxobacteria, a group of Gram-negative, non-firmicutes spore formers, were first assigned to the phylum δ-proteobacteria, but recently reclassified into the newly established phylum *Myxococcota* (Waite *et al*., 2020). *Myxococcus xanthus* is a model organism of myxobacteria. Rod-shaped *M. xanthus* cells can undergo sporulation via two distinct pathways, both resulting in spherical spores (Kroos *et al*., 2025). First, in response to certain chemical signals, such as glycerol and dimethyl sulfoxide (DMSO), individual *M. xanthus* cells can rapidly transition into isolated spherical spores in aqueous environments within hours (Dworkin & Gibson, 1964). Second, millions of cells can aggregate on solid starvation media and develop into spore-filled fruiting bodies, a process that takes a few days to complete (O’Connor & Zusman, 1988). Both sporulation pathways are programmed, tightly controlled processes that involve over 1,000 genes (Muller *et al*., 2010, Munoz-Dorado *et al*., 2019). In contrast to endospore formation, cell division is not involved in either the *M. xanthus* sporulation mechanisms, and *M. xanthus* lacks the homologs of the sporulation genes in *Firmicutes* (Aramayo & Nan, 2022). Thus, sporulating *M. xanthus* must accomplish the rod-to-sphere transition through yet to be discovered PG remodeling mechanisms, which provide invaluable opportunities to investigate PG dynamics (Zhang *et al*., 2021).

Distinct from *Bacilli* and *Clostridia*, *M. xanthus* spores lack cortex PG (Bui *et al*., 2009, Zhang *et al*., 2020, Voelz & Dworkin, 1962). Rather, their resistance is derived from polysaccharide coats (Kottel *et al*., 1975, Perez-Burgos *et al*., 2020, Saidi *et al*., 2021, Muller *et al*., 2012). Sporulating cells that fail to deposit coat polysaccharides on their surfaces produce spherical pseudospores that lack resistance (Wartel *et al*., 2013, Zhang *et al*., 2020, Holkenbrink *et al*., 2014). In a pioneering study, Bui *et al*. did not detect muropeptides in glycerol-induced spores, indicating that *M. xanthus* degrades its PG during rapid sporulation (Bui *et al*., 2009). However, it remains unclear whether starvation-induced spores within fruiting bodies still retain PG and how *M. xanthus* orchestrates programmed morphological changes.

In this report, we used ultra-performance liquid chromatography (UPLC) to demonstrate that *M. xanthus* degrades PG in both sporulation pathways. Through mutagenesis studies, we identified LtgA and LtgB, two lytic transglycosylases (LTGs) essential for these sporulation pathways. Remarkably, LtgB regulates the pace of PG degradation by LtgA during rapid sporulation, preventing the formation of non-resistant pseudospores due to abrupt PG breakdown. This research not only uncovers novel mechanisms of sporulation in non-firmicutes but also highlights the crucial role of cross-regulation between PG hydrolases in maintaining cell integrity.

## Results

### PG is degraded in both spore types

To investigate if starvation-induced spores in *M. xanthus* fruiting bodies retain PG, we broke the fruiting bodies after 120 h of starvation and purified saculli from spores. For comparison, we also purified the sacculi from vegetative cells and glycerol-induced spores. The purification procedure yields only sedimentable PG, as all soluble fragments are removed during the washing steps. Consequently, any PG fragments solubilized by LTG activity during sporulation would be lost at this stage, and the muropeptides we detect derive from the more highly crosslinked material that survives the procedure. The resulting sacculi were then digested with muramidase, and the solubilized muropeptides were analyzed by UPLC (see Materials and Methods). The two spore types showed similar profiles of a discernible presence of muropeptides that resembled those found in vegetative cells, albeit in significantly reduced quantities (***Figure 1***). Importantly, the most abundant muropeptides identified in the vegetative cells, MurNAc-tetrapeptide monomer (M4) and MurNAc-tetrapeptide dimer (D44), were nearly absent in both spore types (***Figure 1***). Such residual muropeptides in both spore types are inadequate for forming continuous PG layers (Bui *et al*., 2009). This observation indicates that spore development in both pathways involves the breakdown of the vegetative PG cell wall.

**Figure 1.**
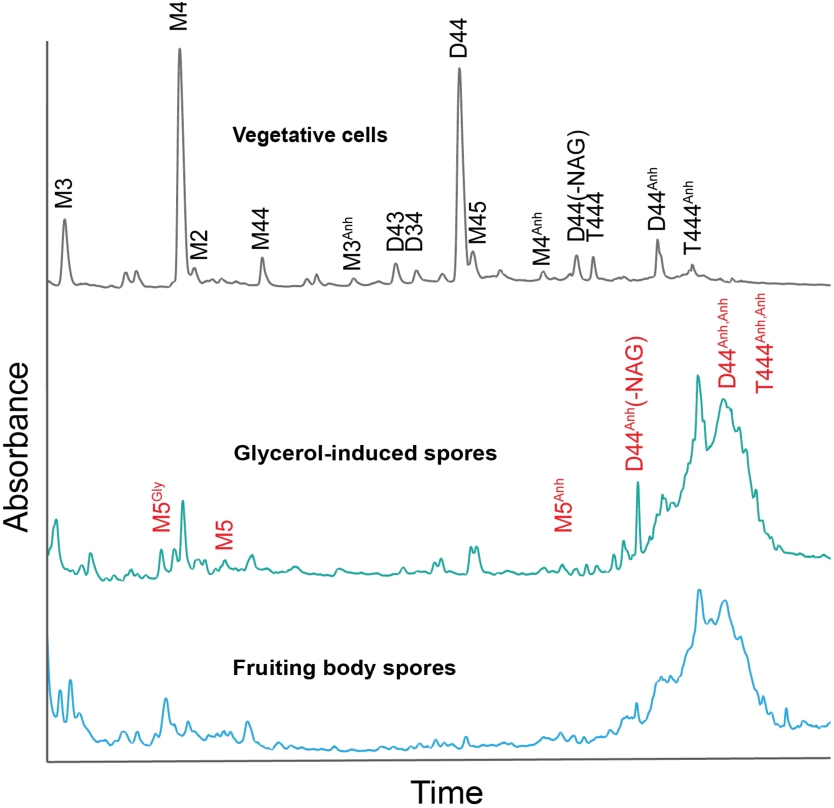
Mature *M. xanthus* spores induced by either glycerol or starvation do not contain significant muropeptides. UPLC muropeptides profiles of vegetative cells, glycerol- and starvation-induced spores indicate that the major muropeptides species, M4 and D44, in vegetative cells are diminished in both spore types, while spores are enriched in anhydro-muropeptides (Anh). Only muropeptides that could be confidently identified by MS/MS analysis are labelled. The characteristic peaks are labeled as follows: M, monomeric muropeptide (uncrosslinked); D, dimeric muropeptide (crosslink connecting two muropeptides), T, trimeric muropeptide (crosslink connecting three muropeptides). Numbers refer to the status of the peptide side chain (3, tripeptide; 4, tetrapeptide). Red characters mark the muropeptides only detected in spores.

Despite the overall decline in identified muropeptides, anhydro-muropeptides — a minor component of vegetative PG — were enriched in both spore types (***Figure 1***). Especially, anhydro-MurNAc-tetrapeptide trimer (T444^Anh^) that was only detected in trace amounts in vegetative cells, became a prominent muropeptide in both spore types (***Figure 1***). Moreover, new anhydro-muropeptides, including anhydro, anhydro-MurNAc-tetrapeptide dimer (D44^Anh,Anh^), anhydro, anhydro-MurNAc-tetrapeptide trimer (T444^Anh,Anh^), and an anhydrodimer without a NAG (D44^Anh^(-NAG)), were only present in spores (***Figure 1***). The identified muropeptides, however, represent only a part of the material released by muramidase from spores: a substantial, as a late-eluting portion of the chromatogram could not be assigned, and its composition remains unknown. Because anhydro-muropeptides are the signature products of LTGs (Dik *et al*., 2017, Williams *et al*., 2018), their abundance in spores indicates that certain LTGs may play important roles in *M. xanthus* sporulation.

### *M. xanthus* sporulation requires two LTGs

The genome of *M. xanthus* encodes 14 putative LTGs (Aramayo & Nan, 2022, Ramirez Carbo *et al*., 2024). We imaged the cells of the 14 knockout mutants (Ramirez Carbo *et al*., 2024) after 6 h of glycerol induction and identified two mutants that displayed abnormality in sporulation (***Figure 2*, *Figure 2* *– figure supplement 1***). We then imaged the wild-type and mutant cells at different time points after glycerol induction and used their length/width (L/W) ratios to monitor potential defects in the sporulation process. Wild-type cells initiated sporulation within 1 h of glycerol induction, which was evidenced by the increase of cell width and decrease of cell length (***Figure 2A***). Consistent with the change in cell morphology, their L/W ratios decreased continuously and stabilized after 2 h of induction, when sporulation completed. In contrast, the cells that carry the deletion of K1515_20820 (MXAN_RS16290) (Aramayo & Nan, 2022) were able to shorten cell length slightly but retained rod shape after prolonged induction (***Figure 2B*, *Figure 2* *– source data 1***).

**Figure 2.**
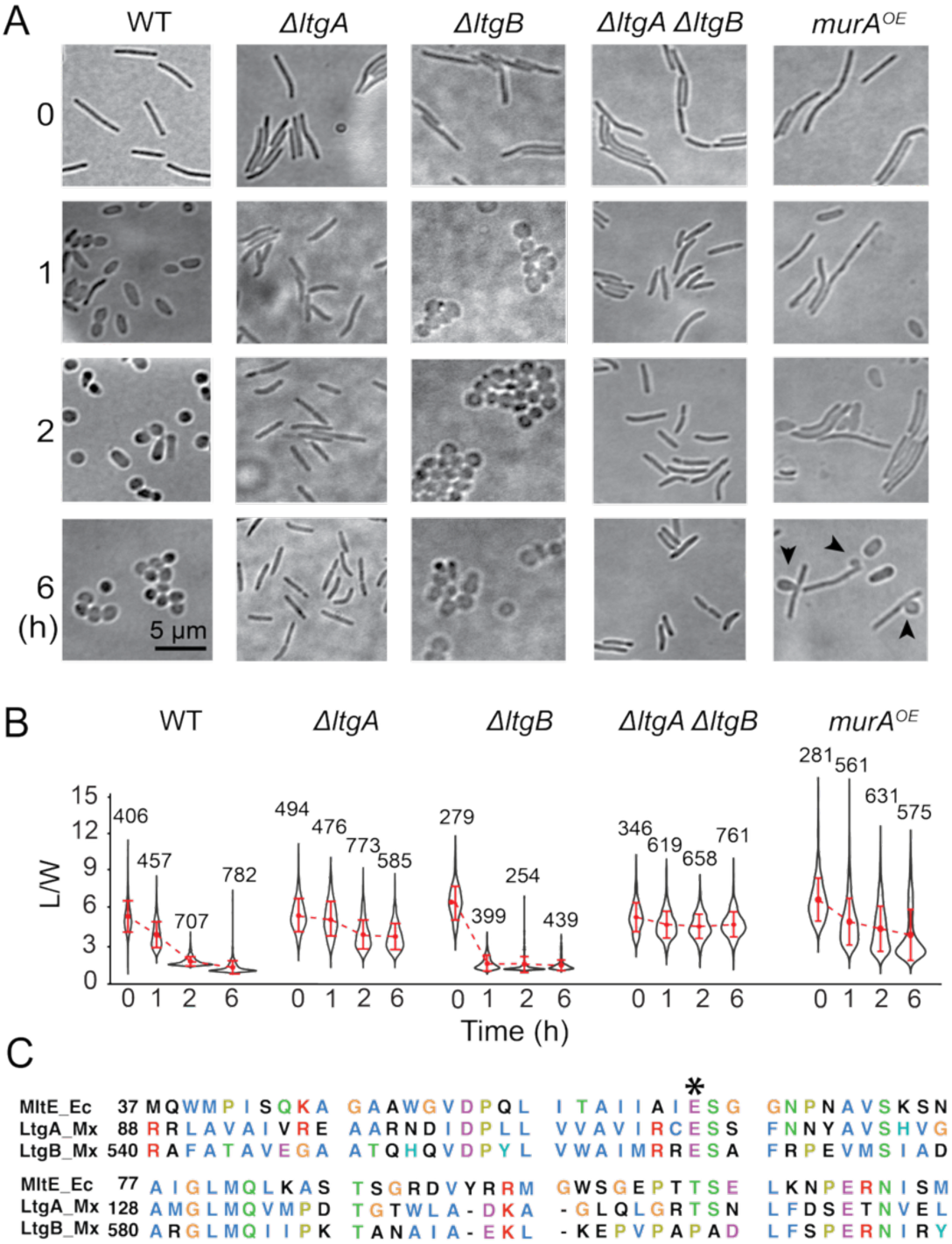
LtgA and LtgB are required for glycerol-induced sporulation, whereas MurA overexpression inhibit sporulation. **A)** Bright field images of cells at different time points after glycerol-induction. Black arrows point to lysing cells. **B)** Quantitative analysis of glycerol-induced sporulation using the length/width ratio (L/W) of cells. Whiskers indicate the 25^th^ - 75^th^ percentiles and red dots the median. The total number of cells analyzed is shown on top of each plot. **C)** LtgA and LtgB are homologous to *E. coli* MltE. The asterisk marks the conserved active site. Other putative LTGs do not affect either sporulation pathway significantly, which is shown in Figure 2 ***– figure supplement 1***. **Source data 1.** Cell morphology measurements for Figure 2B. **Figure supplement 1.** Besides LtgA and LtgB, other putative LTGs do not affect either sporulation pathway significantly. **Figure supplement 2.** In the absence of vanillate, *P_van_*-driven MurA expression does not affect glycerol-induced sporulation.

Surprisingly, another mutant that carries the deletion of K1515_17460 (MXAN_RS19615) started to lose rod shape immediately after glycerol induction and became spherical within 1 h (***Figure 2A*, *2B*, *Figure 2* – source data 1**). However, different from the wild-type spores that appeared dark and heterogeneous under differential interference contrast (DIC) microscopy, the spheres formed by this mutant appeared bright and homogenous, similar to the pseudospores from the *ΔaglQS* mutant that lack the motor for depositing spore coat polysaccharides onto cell surfaces (Zhang *et al*., 2020) (***Figure 2A***). To test whether these spheres are real spores, we subjected them to sonication and quantified their survival rate using a Helber bacterial counting chamber. After sonication, while 91 ± 6% (calculated from three independent experiments, n > 1,000) of the wild-type spores appeared intact, only 3 ± 1% of the spheres produced by the K1515_17460 deletion mutant remained. Thus, this mutant indeed formed pseudospores that lacked the resistance against sonication. As the products of K1515_20820 and K1515_17460 show homology to MltE, an LTG encoded by *Escherichia coli emtA* (49.0% and 38.4% sequence identity, respectively) (***Figure 2C***), we named them *ltgA* and *ltgB*, respectively. Both LtgA and LtgB are required for forming resistant spores via the rapid sporulation pathway, albeit playing different roles in the rod-to-sphere transition.

To determine if these LTGs are also required for starvation-induced sporulation, we grew them in rich liquid media and spotted cells on solid starvation (CF) media. After 96 h of incubation, both mutants produced dark fruiting bodies on the agar surface that were comparable to those of the wild-type (***Figure 3A***). To test if these mutants form starvation-induced spores, we incubated cells in static liquid CF medium using polystyrene Petri dishes for 96 h, scraped the cells attached to the dishes, suspended them in 1 ml water, subjected them to sonication, then plated them on solid rich media. After 96 h of incubation, while the wild-type fruiting bodies produced colonies of 3389 ± 1141 cfu/ml (n = 3), the *ΔltgA* and *ΔltgB* cells failed to form colonies after five days of incubation (***Figure 3B*, *Figure 3* *– source data 1***), indicating that both LtgA and LtgB are essential for forming starvation-induced spores.

**Figure 3.**
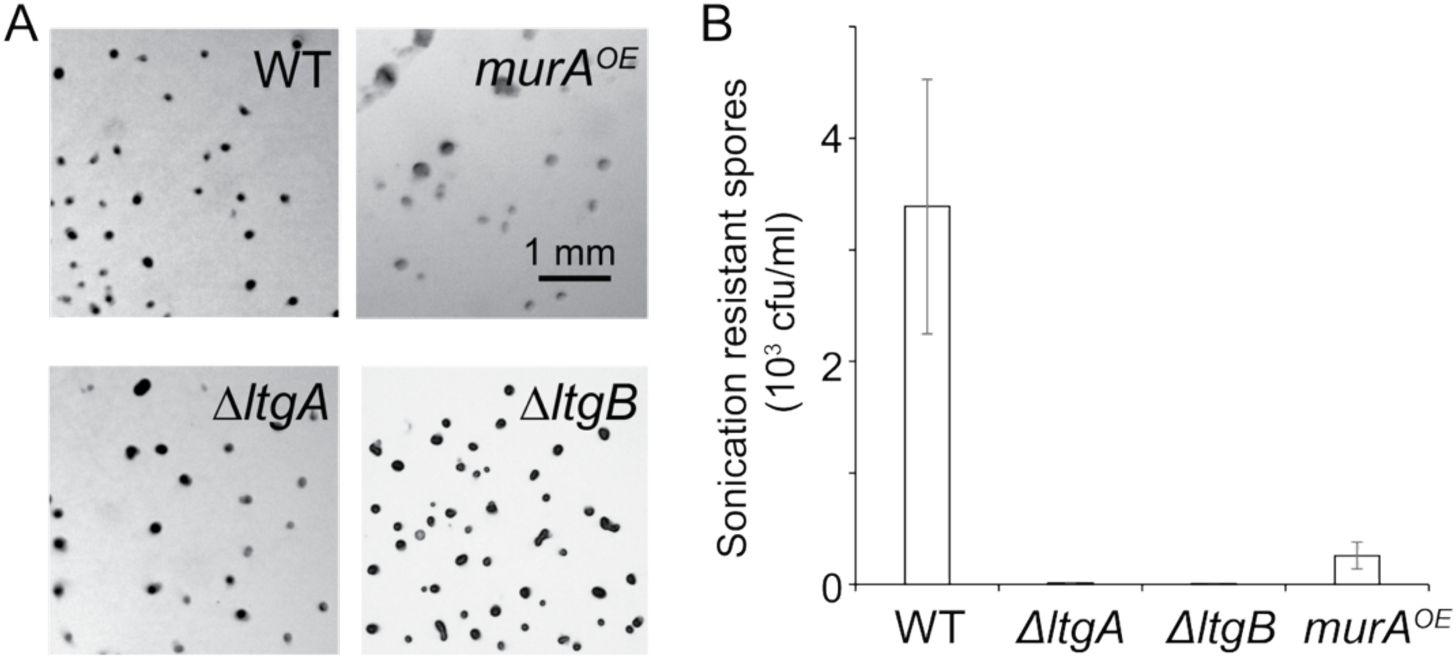
LtgA, LtgB, and MurA regulate starvation-induced sporulation. **A)** The absence of LtgA and LtgB, and the overexpression of MurA do not affect the formation of fruiting body– like aggregates on starvation agar. **B)** Both LtgA and LtgB are essential for forming starvation-induced spores, while MurA overexpression negatively regulates this process. Sonication-resistant spores were counted as colony formation units (cfu) after 96 h of incubation on CYE agar. Data are shown as mean ± standard deviation (n = 3). **Source data 1.** Spore counts for Figure 3B. **Figure supplement 1.** In the absence of vanillate, *P_van_*-driven MurA expression does not affect starvation-induced fruiting body formation.

### LtgA and LtgB play distinct roles in different sporulation pathways

Do LtgA and LtgB degrade PG during sporulation? To answer this question, we purified cell sacculi and used immunofluorescence and an anti-PG serum (de Pedro *et al*., 1997) to visualize the remaining PG. The sacculi of vegetative cells from the wild-type, *ΔltgA*, and *ΔltgB* strains were not visible under DIC microscopy, likely due to their flattened structures minimizing light interference. However, PG from all three strains was readily detected in the fluorescence channel (***Figure 4A***). After 6 h of glycerol-induction, the sacculi of both the wild-type and *ΔltgA* cells remained visible under DIC microscopy (***Figure 4A***), likely due to the deposition of spore coat polysaccharides that sustained unflattened cell structures (Wartel *et al*., 2013, Holkenbrink *et al*., 2014). After the sacculus purification process, only background PG signals were detected in glycerol-induced wild-type spores (***Figure 4A***). In contrast, the *ΔltgA* cells retained PG in substantial quantities (***Figure 4A***). Sacculi from the *ΔltgB* pseudospores still contained PG but showed lower fluorescence intensity (***Figure 4A***). While the remaining PG sacculi of these pseudospores were largely spherical, they lost integrity during purification, with many sacculi displaying irregular shapes in the fluorescence channel (***Figure 4A***). Thus, compared to LtgA, the enzymatic activity of LtgB plays a more limited role in PG degradation during chemical-induced sporulation. Collectively, LtgB appears to be a pace-keeper that prevents abrupt PG degradation. Different from the induced wild-type and *ΔltgA* cells, sacculi from the *ΔltgB* pseudospores were undetectable under DIC microscopy (***Figure 4A***), reflecting their flattened shapes. This phenotype suggests that further acceleration of the morphological transitions associated with rapid sporulation may impede the formation of a rigid polysaccharide coat.

**Figure 4.**
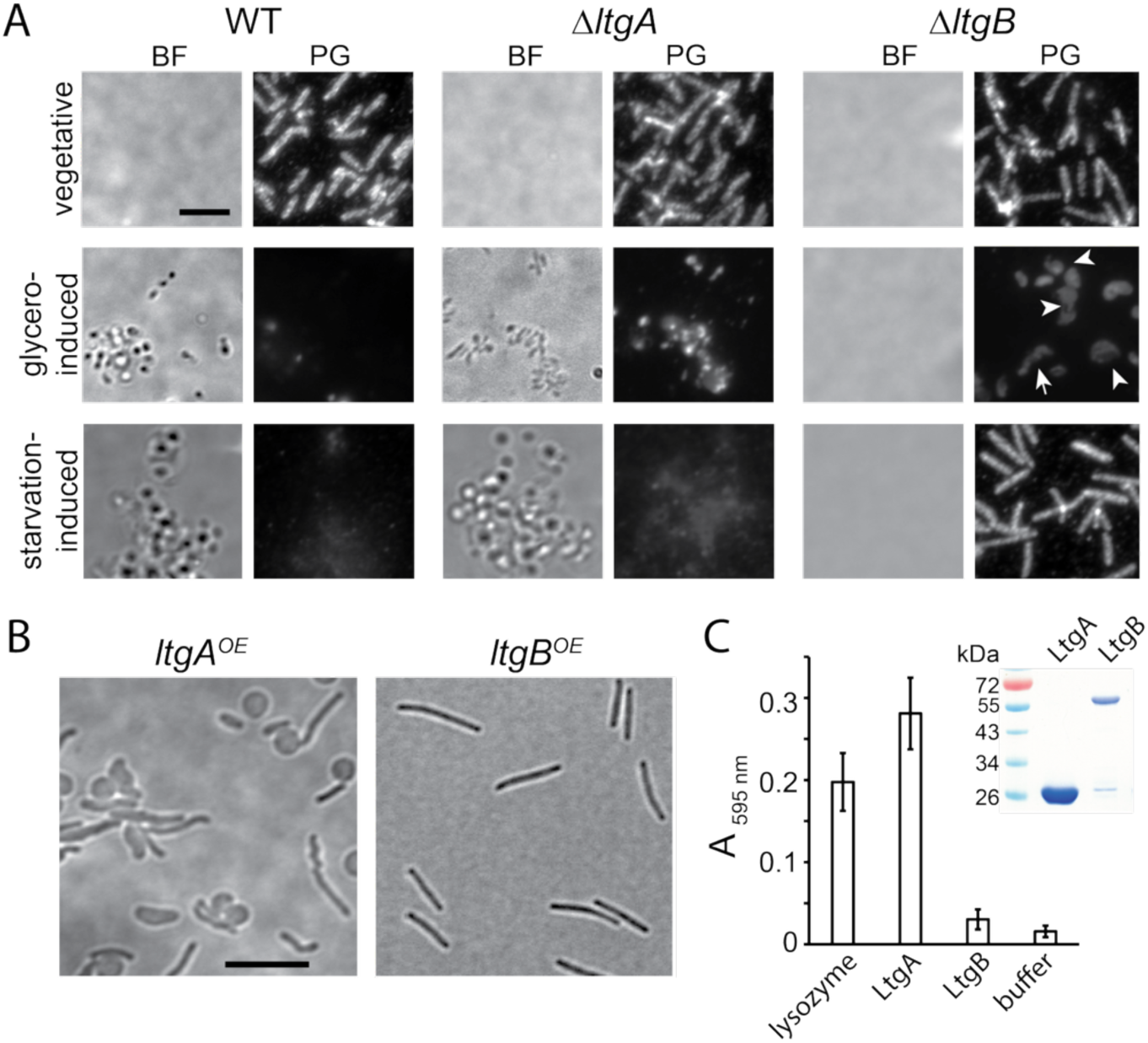
LtgA and LtgB play distinct roles in the two sporulation pathways. **A)** While LtgA is required for PG degradation during glycerol-induced sporulation, LtgB is the major LTG for forming starvation-induced spores. PG was detected using an anti-PG serum and a fluorescence-conjugated secondary antibody. PG sacculi were purified from cells after 6 h and 120 h of glycerol-induced and starvation-induced sporulation, respectively. Flattened sacculi are not visible under bright field microscopy. White arrows point to the sacculi in irregular shapes. BF, bright field. **B)** The overexpression of LtgA, but not LtgB, collapses rod-shape in vegetative cells. **C)** Purified LtgA and LtgB solubilize dye-labeled PG sacculi at different rates. Lysozyme and buffer serve as the positive and negative controls, respectively. Absorption at 595 nm was measured after 18 h incubation at 25 ⁰C. Data are presented as mean values ± SD from three technical replicates. The inset shows purified LtgA and LtgB in a Coomassie stained gel. Scale bars, 5 μm. **Source data 1.** Absorption at 595 nm in the Remazol brilliant (RBB) assay for Figure 4C. **Figure supplement 1.** Vanillate-induced expression of LtgA-PAmCherry and LtgB-PAmCherry.

To determine the roles of LtgA and LtgB in slow sporulation, we first scraped the wild-type fruiting bodies from CF agar surface (***Figure 2C***), dispersed spores by sonication, purified their sacculi, and visualized PG using immunofluorescence. Spherical spores from the wild-type cells were visible under DIC microscope, indicating that they retained unflattened shapes. These fruiting body spores lacked PG-specific fluorescence (***Figure 4A***), consistent with their markedly reduced PG content (***Figure 1***). However, we cannot exclude the possibility that the polysaccharide spore coats (Voelz & Dworkin, 1962) hinder antibody access to PG. Second, we investigated if *ΔltgA* and *ΔltgB* cells degraded PG during starvation-induced sporulation. Because the aggregates formed by *ΔltgA* and *ΔltgB* cells on CF agar did not contain mature spores, we scrapped cell aggregates from agar surface and purified their sacculi without sonication. The *ΔltgA* cells formed spherical, spore-like cells void of PG (***Figure 4A***), indicating that while LtgB is sufficient for PG degradation during slow sporulation, LtgA is only required for spore maturation. In contrast, most of the *ΔltgB* cells were still rod-shaped, indistinguishable from vegetative ones (***Figure 4A***). These observations indicate that LtgB is an essential LTG for PG degradation during slow sporulation.

The two sporulation pathways vary greatly in duration: glycerol-induced rapid sporulation is completed within four hours, while starvation-induced slow sporulation unfolds over several days. Based on this distinction, we hypothesized that LtgA and LtgB could degrade PG at different rates. We used a vanillate-inducible promoter (Iniesta *et al*., 2012) to overexpress LtgA and LtgB as a merodiploids. Upon induction with 200 µM vanillate, cells overproducing LtgA exhibited heterogeneous morphology, likely resulting from variations in LtgA expression or differences in cell cycle stages within the population. Many cells lost rod shape even in the absence of glycerol, indicating that these cells over-degraded their PG by LtgA (***Figure 4B***). In contrast, the overexpression of LtgB using the same method did not affect the morphology of vegetative cells (***Figure 4B***). To test if LtgB overexpression was induced by vanillate, we added a photo-activatable mCherry (PAmCherry) tag to the overexpression constructs of LtgA and LtgB. As shown in ***Figure 4* *– figure supplement 1***, in the presence of vanillate, the expression of both LTGs increased significantly. These findings suggest that LtgB exhibits very low LTG activity in vegetative cells, as overexpression of the protein fails to induce substantial PG degradation.

To further investigate the activities of these LTGs, we expressed the periplasmic domains of wild-type LtgA (amino acids 21 – 243) and LtgB (amino acids 26 – 710) in *E. coli* (***Figure 4C*, *Figure 4* *– source data 1***). We purified PG from wild-type *M. xanthus* cells, labeled it with Remazol brilliant blue (RBB), and tested if the purified LTGs hydrolyze labeled PG *in vitro* and release the dye (Jorgenson *et al*., 2014, Ramirez Carbo *et al*., 2024, Uehara *et al*., 2010). Wild-type LtgA solubilized dye-labeled PG, which absorbed light at 595 nm, demonstrating stronger hydrolytic activity than lysozyme, an enzyme that specifically cleaves β-1,4-glycosidic bonds in PG. In contrast, PG incubated with LtgB only showed minimum release of the dye, indicating slow PG hydrolysis (***Figure 4C*, *Figure 4* *– source data 1***). While we cannot rule out the possibility that our purification and reaction conditions were suboptimal for LtgB, both the *in vitro* RBB assay and the *in vivo* phenotype resulting from LtgB overexpression support our hypothesis that during vegetative growth, LtgB is less efficient in PG degradation than LtgA.

### LtgB regulates LtgA during glycerol-induced sporulation

Consistent with its role in PG degradation, in a microarray-based transcriptome analysis, *ltgA* transcription was found to increase about twofold during rapid sporulation (4 h) but remain unchanged during slow sporulation (96 h) in the closely related DK1622 strain (Muller *et al*., 2010). Conversely, *ltgB* expression gradually rises during slow sporulation, reaching 1.8 times the vegetative level at 96 h, while remaining stable during rapid sporulation (Muller *et al*., 2010, Munoz-Dorado *et al*., 2019). However, transcriptomic data alone do not account for the opposing roles of LtgA and LtgB in the rod-to-sphere transition during rapid sporulation. The resemblance in morphology between glycerol-induced *ΔltgB* cells and uninduced cells overproducing LtgA (***Figure 4B***) suggests that the slower-acting LtgB may offset the rapid activity of LtgA during sporulation, allowing sufficient time for spore maturation.

To test if LtgB regulates LtgA, we constructed a *ΔltgA ΔltgB* double deletion strain. Cells from this strain shortened their length slightly but failed to abolish rod shape after prolonged glycerol induction, phenocopying the sporulation defect of the *ΔltgA* strain (***Figure 2A*, 2B, *Figure 2* *– source data 1***). Hence, LtgB is a regulator upstream of LtgA.

### LtgB exhibits a response to glycerol induction earlier than LtgA

Biochemical reactions on PG are unique for the stark size difference between the enzymes and their substrates. While the enzymes are in nanometer scales, their substrates, the PG sacculi, span several micrometers. Under the microscope, PG remains stationary but PG-related enzymes are free to move. Even for *M. xanthus* that moves on surfaces, PG-related enzymes move at least two orders of magnitude faster than the cell/PG (Zhang *et al*., 2023b, Ramirez Carbo *et al*., 2024). Thus, when diffusive enzymes bind to PG, their mobility decreases (Lee *et al*., 2016, Zhang *et al*., 2023b). For instance, DacB, another PG hydrolase, reduces its single-particle mobility in the conditions where its activity is activated (Zhang *et al*., 2023b). By tracking single fluorescently-labeled enzyme particles, we can approximate their PG-binding in different physiological conditions and genetic backgrounds (Ramirez Carbo *et al*., 2024, Zhang *et al*., 2023b, Ramírez Carbó & Nan, 2026). We individually expressed PAmCherry-labeled LtgA and LtgB using their native loci and promoters. Both labeled LTGs accumulated as full-length proteins (***Figure 5* *– figure supplement 1A***) and the PAmCherry tags did not negatively affect either glycerol or starvation-induced sporulation pathways, indicating that these fusion proteins were fully functional (***Figure 5* *– figure supplement 1B, C***). Notably, vegetative cells produce significantly less LtgA than LtgB, consistent with the RNAseq data in which *ltgA* transcription was below the detection limit (Munoz-Dorado *et al*., 2019) (***Figure 5* *– figure supplement 1A***).

**Figure 5.**
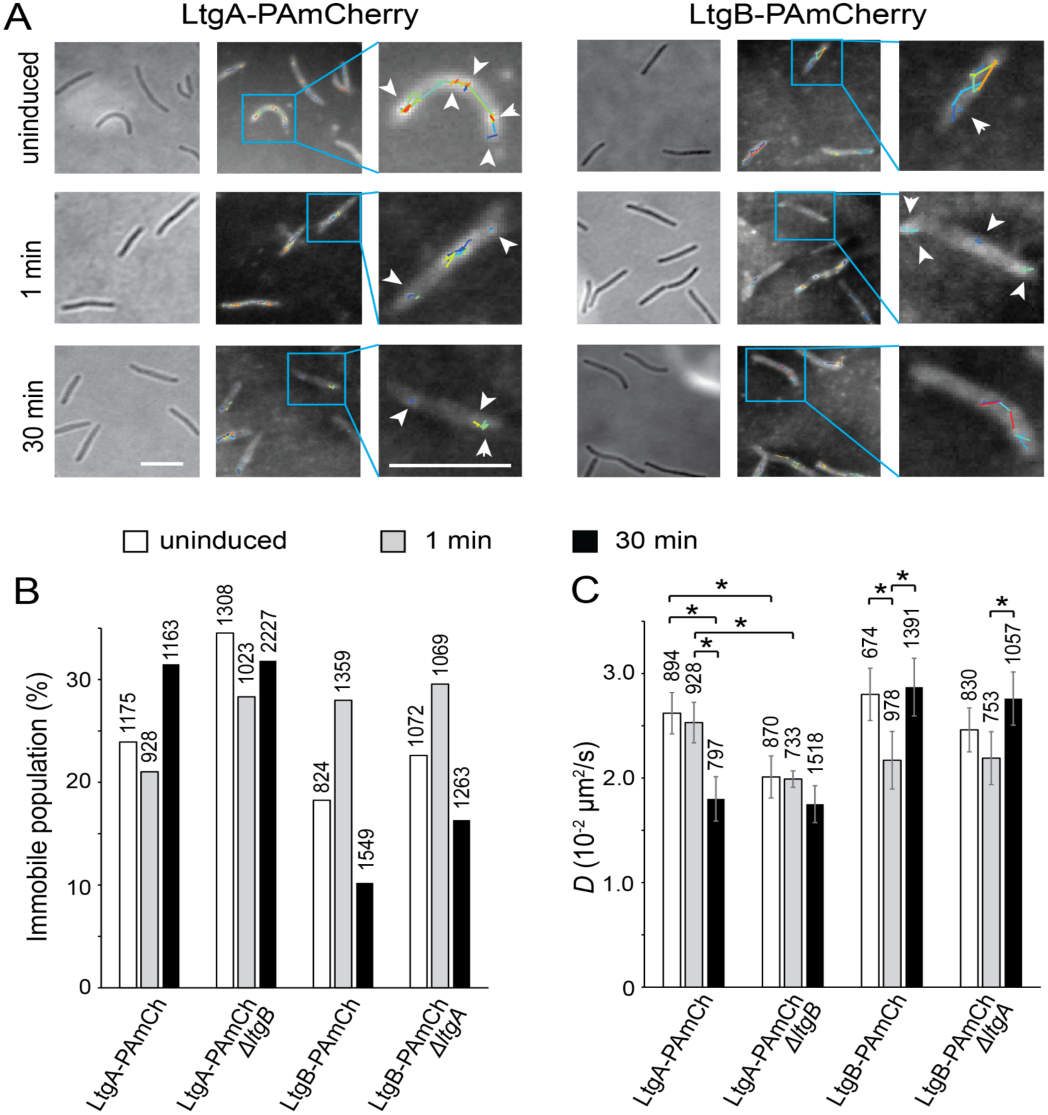
LtgB regulates the PG-binding of LtgA during glycerol-induced sporulation. **A)** LtgB exhibits a response to glycerol induction earlier than LtgA. Representative trajectories of LtgA and LtgB before (uninduced) and after (1 min and 30 min) glycerol induction. The overall distribution of both LTGs is displayed using the composite of 100 consecutive frames taken at 100-ms intervals. Single-particle trajectories of PAmCherry were generated from the same frames. Individual trajectories are distinguished by colors. Scale bars, 5 μm. **B** and **C)** the absence of LtgB reduces the diffusion of LtgA, which is reflected in the increase of immobile population (**B**) and the decrease in diffusion coefficient (*D*) (**C**), and these effects are especially prominent in the cells before (uninduced) and immediately after (1 min) glycerol induction. For each protein and condition, particles were identified from at least 100 cells and three independent experiments. The total number of particles analyzed is shown on top of each plot. Error bars were the standard derivation of 1,000 bootstrap samples and * indicates a difference of > 0.005. **Source data 1.** Diffusion coefficient data for Figure 5C. **Figure supplement 1.** PAmCherry-labeled LtgA and LtgB express as full-length and fully functional proteins.

We used a 405-nm excitation laser (0.3 kW/cm^2^, 0.1 s) to activate the fluorescence of a few labeled LTG particles randomly in each cell and quantified their localization using a 561-nm laser at 10 Hz using single particle tracking photo-activated localization microscopy (sptPALM, see Materials and Methods) (Fu *et al*., 2018, Nan *et al*., 2015, Nan *et al*., 2013, Ramírez Carbó & Nan, 2026). As free PAmCherry particles diffuse extremely fast in the cytoplasm, entering and exiting the focal plane frequently, they usually appear as blurry objects that cannot be followed at 10 Hz close to the cell surface (Fu *et al*., 2018, Zhang *et al*., 2023b). For this reason, the noise from free PAmCherry due to potential protein degradation was negligible.

Single-particles of PAmCherry-labeled LtgA and LtgB can be categorized into two populations, immobile and mobile. The immobile particles remained within a single pixel (160 nm × 160 nm) before photobleach, and the mobile ones displayed typical diffusion (***Figure 5A*, *Figure 5* *– source data 1***). For the particles that switched between mobile and immobile states, our algorithm categorized them as mobile and calculated their diffusion coefficients (*D*) from their entire trajectories that contained both mobile and immobile segments. Hence, binding to PG not only increases the immobile population of the enzyme particles but also decreases the *D* of the mobile particles. In vegetative cells where large-scale PG degradation does not occur, 23.9% (n = 1175) and 18.2% (n = 824) of LtgA and LtgB particles were immobile, respectively (***Figure 5B*, *Figure 5* *– source data 1***). *D* values of the mobile population were (2.62 ± 0.20) × 10^-2^ µm^2^/s (n = 894) for LtgA-PAmCherry and (2.80 ± 0.25) × 10^-2^ µm^2^/s (n = 674) for LtgB-PAmCherry (***Figure 5C*, *Figure 5* *– source data 1***).

We then determined how the dynamics of both LTGs respond to glycerol induction in the rapid sporulation pathway. Immediately (1 min) after adding glycerol, the dynamics of LtgA remained little changed, when 21.0% (n = 928) of particles were immobile and the *D* of the mobile ones was (2.53 ± 0.19) × 10^-2^ µm^2^/s (n = 733) (***Figure 5B*, C, *Figure 5* *– source data 1***). In contrast, the mobility of LtgB decreased significantly, with the immobile population increased to 28.0% (n = 1359) and *D* of the mobile population decreased to (2.17 ± 0.28) × 10^-2^ µm^2^/s (n = 978) (***Figure 5B*, C, *Figure 5* *– source data 1***). Therefore, these results suggest that upon glycerol induction, LtgB rapidly enhances its binding to PG.

We then chose 30 min after glycerol induction as a time point for rapid PG degradation, which is reflected by the dramatic decrease of L/W during the first hour of sporulation (***Figure 2A*, *B*, *Figure 2* *– source data 1***). Compared to its inert response 1 min after glycerol induction, the mobility of LtgA particles decreased significantly at 30 min, when the immobile population increased to 31.5% (n = 1163) and *D* of the mobile population decreased to (1.80 ± 0.21) × 10^-2^ µm^2^/s (n = 797) (***Figure 5B*, *C*, *Figure 5* *– source data 1***). The reduced mobility of LtgA aligns with its function as the primary LTG in glycerol-induced sporulation. Strikingly different from LtgA, as sporulation advanced to the 30-min mark, LtgB’s mobility returned to its pre-induction level, with the immobile population decreased to 10.2% (n = 1549) and *D* of the mobile population increased to (2.87 ± 0.28) × 10^-2^ µm^2^/s (n = 1391) (***Figure 5B***, ***C*, *Figure 5* *– source data 1***). Therefore, the PG-binding of LtgB shows a negative correlation with PG degradation. Taken together, LtgA and LtgB display opposite responses to glycerol induction, in which LtgB rapidly binds to PG before yielding to LtgA, whose PG-binding is concurrent with the rod-to-sphere transition.

### LtgB blocks LtgA from binding PG in the early stage of glycerol-induced sporulation

Does LtgB’s early PG-binding suppress PG degradation by LtgA? To answer this question, we expressed LtgA-PAmCherry using the native *ltgA* locus and promoter in the *ΔltgB* background. In vegetative cells, the absence of LtgB significantly reduced the mobility of single LtgA-PAmCherry particles, suggesting that LtgB does affect LtgA’s binding to the PG (***Figure 5B***, ***C*, *Figure 5* *– source data 1***). Similarly, we expressed LtgB-PAmCherry using the native *ltgB* locus and promoter in the *ΔltgA* background. In contrast, the mobility of single LtgB-PAmCherry particles only decreased slightly in the absence of LtgA (***Figure 5B***, ***C*, *Figure 5* *– source data 1***). These results suggest that while LtgB significantly reduces LtgA’s binding to PG, probably due to its higher expression level, LtgA has only a modest impact on LtgB’s ability to bind PG.

Strikingly different from its slow response to glycerol in wild-type cells, LtgA increased its PG-binding immediately (1 min) after glycerol induction in the *ΔltgB* background, confirming that LtgB may compete with LtgA for access to PG in the early stage of sporulation (***Figure 5B***, ***C*, *Figure 5* *– source data 1***). In contrast, at 30 min of induction, a time point when LtgB dissociated from PG, its absence no longer affected LtgA’s PG-binding (***Figure 5B***, ***C*, *Figure 5* *– source data 1***). Similar to the observation in wild-type cells, the lack of LtgA did not affect the PG-binding of LtgB at either 1 min or 30 min after glycerol induction (***Figure 5B***, ***C*, *Figure 5* *– source data 1***). In summary, the sequential PG-binding by these two LTGs controls the pace of PG-degradation during glycerol-induced sporulation. LtgB, a slow LTG, regulates the progression of rapid sporulation by preventing LtgA, a fast LTG, from excessively degrading PG, thereby ensuring spore maturation.

### Upregulated PG synthesis negatively affects PG degradation

Since LtgA is less mobile in vegetative *ΔltgB* cells, i.e. strongly binds to PG (***Figure 5B***, ***C*, *Figure 5* *– source data 1***), why do these cells retain their rod shape rather than losing it spontaneously before glycerol induction? In addition to PG-binding, LtgA may require additional stimuli to initiate PG degradation. We hypothesized that, similar to certain antibiotics targeting PG synthases, which induce wild-type cells to degrade PG and form pseudospores in rich liquid media (Zhang *et al*., 2023b, O’Connor & Zusman, 1997), glycerol may alter cellular metabolism, leading to reduced PG synthesis. If this is the case, cells with upregulated PG synthesis should remain rods even in the presence of glycerol.

To test our hypothesis, we used the vanillate-inducible promoter (Iniesta *et al*., 2012) to overexpress *murA* as a merodiploid in the wild-type background. Because MurA catalyzes the first committed step of PG synthesis that produces UDP-MurNAc (Egan *et al*., 2020, Rohs & Bernhardt, 2021), elevated MurA levels are expected to channel more cellular resources toward PG synthesis. Overproduction of MurA in the presence of 200 µM vanillate resulted in a heterogeneous cell population, with normal cells coexisting alongside elongated ones (***Figure 2A***, ***B*, *Figure 2* *– source data 1***). Similar to the cells that overexpressed LtgA (***Figure 4B***), this heterogeneity likely reflects variable *murA* induction or the unsynchronized growth stages within the population. Overall, excessive MurA increased the average length of vegetative cells by 19.6%. Cells overproducing MurA also displayed heterogeneity during glycerol-induced sporulation. While some cells transitioned into spheres, many retained their rod shape even after 6 h of induction (***Figure 2A***, ***B*, *Figure 2* *– source data 1***). Notably, sporadic lysis of rod-shaped cells began 2 h post-induction and became increasingly frequent with continued incubation (***Figure 2A***). Potentially, elevated muropeptide production may lead to the accumulation of toxic intermediates that the relatively insufficient LTGs failed to degrade (Weaver *et al*., 2022). Nevertheless, the capacity of MurA overexpression to enable certain cells to circumvent glycerol-induced sporulation implies that PG production acts as a regulatory cue for PG degradation.

To test if excessive MurA can also reduce PG degradation during slow sporulation, we induced MurA overexpression on solid CF agar containing 100 µM vanillate. After 96 h of incubation, these cells developed fruiting body-like aggregates (***Figure 3A***). To test if such fruiting bodies contained mature spores, we scraped them from submerged CF medium and tested the germination rate of spores on CYE agar after sonication. The fruiting bodies formed by cells overexpressing MurA yielded 7.6% of the colonies produced by an equivalent number of wild-type cells, indicating that upregulated PG synthesis also prevents slow sporulation. In contrast, cells grown without vanillate progressed normally through glycerol-induced sporulation and starvation-induced fruiting body formation, showing no differences compared to wild-type cells (***Figure 3B* *– Figure supplement 2***). Therefore, the sporulation of *M. xanthus* is sensitive to PG synthesis and increased PG synthesis balances PG degradation in both sporulation pathways.

## Discussion

Our findings demonstrate that *M. xanthus*, a non-firmicute bacterium, relies on PG degradation to change cell shape during sporulation. This mechanism presents a significant challenge for sporulating cells: preserving their structural integrity while simultaneously dismantling the primary framework that upholds it. Our findings revealed that besides regulating the production of LTGs, cells could control the pace of PG degradation in a rapid sporulation pathway through the regulation between two LTGs and hence allow spore maturation.

Studying the regulation between two LTGs is especially challenging both *in vivo* and *in vitro* for a few reasons. First, it is difficult to monitor their instantaneous activities *in vivo*. Secondly, muropeptide analysis may not distinguish their roles because their products bear the same signature, anhydro-MurNAc. Thirdly, some enzymes require special substrates or activators, which would be difficult to study *in vitro*. To overcome these technical hurdles, we leveraged single-particle mobility to quantify the PG-binding of these enzymes. The simultaneous occurrence of reduced LtgA mobility and PG degradation during glycerol-induced sporulation indicates that the molecular dynamics of LtgA accurately mirrors its enzymatic activity. Such correlation between decreased particle mobility and increased enzymatic activity applies to many other PG-related enzymes, including multiple PG polymerases in *E. coli* and the endopeptidase DacB in *M. xanthus* (Lee *et al*., 2016, Zhang *et al*., 2023b, Yang *et al*., 2021). Because the size difference between PG and its related enzymes is a conserved feature in all bacteria, our method using single-particle tracking to quantify PG-binding can be applied in many organisms.

We discovered that LtgB, a slower-acting LTG, limits the access of the faster LtgA to the PG during the early phase of rapid sporulation, thereby regulating the rate of its degradation. Then how does LtgA breach the blockage of LtgB and gain access to PG as sporulation progresses? Because the overexpression of LtgA is sufficient to damage PG in uninduced cells (***Figure 4B***), we propose that the relative abundance of these two LTGs plays a critical role in determining the fate during fast sporulation. While LtgB nearly saturates PG binding sites in vegetative cells where few LtgA are produced, LtgA—upregulated after glycerol induction (Muller *et al*., 2010) —gradually outcompetes LtgB and thereby causes LtgB to disassociate from PG.

Although glycerol has been known for decades to trigger sporulation in *M. xanthus*, the underlying mechanism remains elusive. Notably, glycerol was recently found to disrupt peptidoglycan (PG) synthesis in *B. subtilis*; in contrast, this effect inhibits *B. subtilis* sporulation, which depends on enhanced PG synthesis (Updegrove et al., 2026). While elevated glycerol concentrations are associated with reduced PG synthesis in both species, whether and how glycerol affects PG synthesis in *M. xanthus* remains unclear. Furthermore, it is unknown whether this apparent correlation reflects a shared evolutionary origin or a common mechanistic basis. Nevertheless, PG dynamics must play critical roles in bacterial sporulation

As increased PG synthesis enables many *M. xanthus* cells to bypass both sporulation pathways, it is reasonable to propose that diminished PG synthesis initiates PG degradation by LTGs. While functional coordination between PG synthesis and hydrolysis has been speculated for over 50 years (Koch, 1985, Koch, 1990), its mechanisms only came to light recently. Some endopeptidases are proposed to serve as the space-makers for PG synthases (Dorr *et al*., 2013, Singh *et al*., 2012). In *M. xanthus*, we reported that moenomycin, an antibiotic that specifically binds to the GTase domains of class A penicillin-binding proteins (aPBPs) (Sung *et al*., 2009, Lovering *et al*., 2007), specifically activates a PG endopeptidase DacB through PBP1a2, an aPBP (Zhang *et al*., 2023b). Because moenomycin mimics a growing glycan strand in the GTase domains of aPBPs, it actually locks aPBPs in their glycan-charged conformations (Sung *et al*., 2009, Lovering *et al*., 2007). Thus, similar to the moenomycin-bound form, glycan-charged PBP1a2 in physiological conditions recruits DacB to the PG assembly sites, which in turn, generates openings in the existing PG network as the crosslink sites for the TPase activity of PBP1a2. Such “make-before-break” mechanism (Koch & Doyle, 1985) could prevent unneeded hydrolysis and thus maintain PG integrity.

Compared to the well-characterized space-making functions of endopeptidases, the roles of LTGs, beyond PG recycling, remain ill-defined. Even less is known on their potential coordination with PG synthases (Weaver *et al*., 2023). Because LTGs from bacterial and phage origins are inhibited by L,D-crosslinks, they may functionally connect to PBPs that form D,D-crosslinks (Alvarez *et al*., 2024). Additionally, some LTGs, such as MltD and MltG in *E. coli* and MltG in *Vibrio cholerae*, prefer to degrade nascent, uncrosslinked glycan strands (Kaul *et al*., 2024, Weaver *et al*., 2022, Bohrhunter *et al*., 2021). Our recent report indicates that AgmT, an *M. xanthus* LTG homologous to *E. coli* MltG, detoxifies uncrosslinked glycan strands under antibiotic stress and modifies crosslinked PG scaffolds to attach a motility machinery to PG (Ramirez Carbo *et al*., 2024). Then how does reduced PG synthesis activate LtgA in *M. xanthus*? Although MltE, the closest homolog of LtgA in *E. coli*, degrades glycan strands regardless of their crosslinking status or peptide content, it shows a marked preference for uncrosslinked substrates (Dik *et al*., 2017, Fibriansah *et al*., 2012). Similar to AgmT and MltE, LtgA might also bind to both crosslinked and uncrosslinked glycan strands. Consistent with its high mobility, LtgA may coordinate with diffusive PG polymerases and cleave the uncrosslinked glycan strands. During rapid sporulation, as nascent glycan strands decline due to muropeptide depletion, LtgA increasingly binds to the crosslinked PG scaffold, which is evidenced by its reduced mobility. This binding intensifies further as PG degradation continues, establishing a positive feedback loop. In this case, LtgB serves as a brake that dampens this positive feedback, moderates PG degradation, and thus allows sporulating cells to maintain integrity by assembling polysaccharide coats. Such cross-regulation between PG hydrolases, which is currently under investigation, provides yet another mechanism by which bacteria dynamically maintain their PG cell walls.

## Materials and Methods

### Bacterial strains and growth conditions

Vegetative *M. xanthus* cells were grown in liquid CYE medium (10 mM MOPS pH 7.6, 1% (w/v) Bacto™ casitone (BD Biosciences), 0.5% yeast extract and 8 mM MgSO_4_) at 32 °C, in 125-ml flasks with vigorous shaking, or on CYE plates that contains 1.5% agar. We used strain DZ2 as the wild-type *M. xanthus* strain (Campos & Zusman, 1975). Knock-out mutants were constructed by electroporating DZ2 cells with 4 µg of plasmid DNA. Transformed cells were plated on CYE plates supplemented with 100 mg/ml sodium kanamycin sulfate or 10 mg/ml tetracycline. The strains and plasmids used in this study are listed in **Table 1**.

**Table 1.**
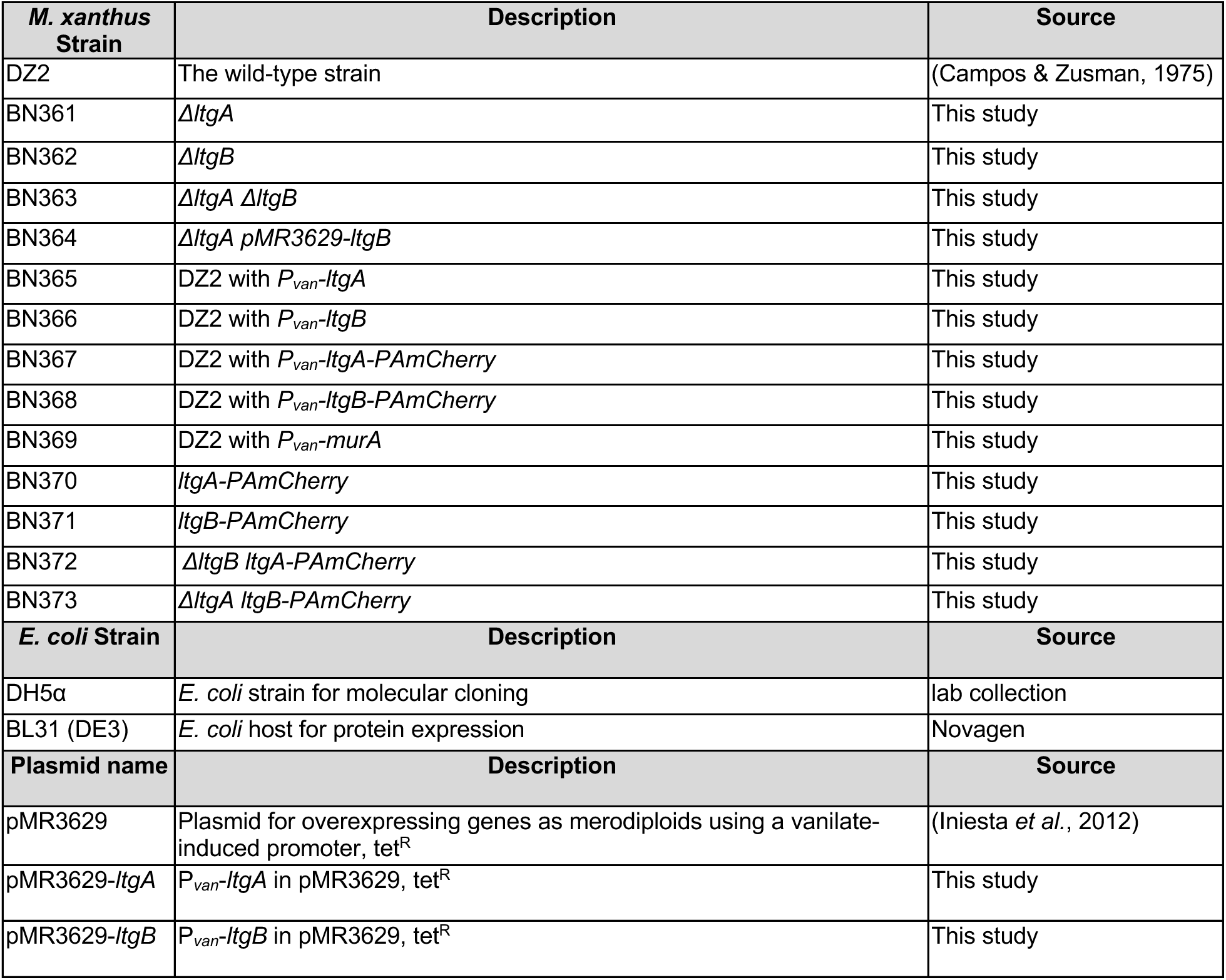
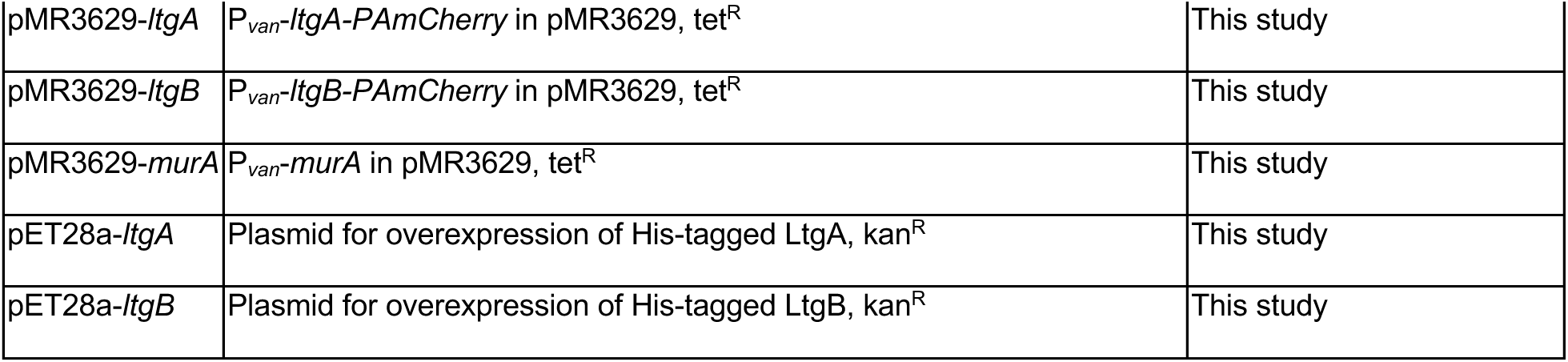
Strains and plasmids used in this study.

| *M. xanthus* Strain | Description | Source |
| --- | --- | --- |
| DZ2 | The wild-type strain | (Campos & Zusman, 1975) |
| BN361 | Δ*ItgA* | This study |
| BN362 | Δ*ItgB* | This study |
| BN363 | Δ*ItgA* Δ*ItgB* | This study |
| BN364 | Δ*ItgA* pMR3629-*ItgB* | This study |
| BN365 | DZ2 with *P<sub>van</sub>-ItgA* | This study |
| BN366 | DZ2 with *P<sub>van</sub>-ItgB* | This study |
| BN367 | DZ2 with *P<sub>van</sub>-ItgA-PAmCherry* | This study |
| BN368 | DZ2 with *P<sub>van</sub>-ItgB-PAmCherry* | This study |
| BN369 | DZ2 with *P<sub>van</sub>-murA* | This study |
| BN370 | *ItgA-PAmCherry* | This study |
| BN371 | *ItgB-PAmCherry* | This study |
| BN372 | Δ*ItgB ItgA-PAmCherry* | This study |
| BN373 | Δ*ItgA ItgB-PAmCherry* | This study |
| *E. coli* Strain | Description | Source |
| DH5α | *E. coli* strain for molecular cloning | lab collection |
| BL31 (DE3) | *E. coli* host for protein expression | Novagen |
| Plasmid name | Description | Source |
| pMR3629 | Plasmid for overexpressing genes as merodiploids using a vanilate-induced promoter, tet<sup>R</sup> | (Iniesta *et al.*, 2012) |
| pMR3629-*ItgA* | *P<sub>van</sub>-ItgA* in pMR3629, tet<sup>R</sup> | This study |
| pMR3629-*ItgB* | *P<sub>van</sub>-ItgB* in pMR3629, tet<sup>R</sup> | This study |
| pMR3629-*ltgA* | P<sub>van</sub>-*ltgA*-PAmCherry in pMR3629, tet<sup>R</sup> | This study |
| pMR3629-*ltgB* | P<sub>van</sub>-*ltgB*-PAmCherry in pMR3629, tet<sup>R</sup> | This study |
| pMR3629-*murA* | P<sub>van</sub>-*murA* in pMR3629, tet<sup>R</sup> | This study |
| pET28a-*ltgA* | Plasmid for overexpression of His-tagged LtgA, kan<sup>R</sup> | This study |
| pET28a-*ltgB* | Plasmid for overexpression of His-tagged LtgB, kan<sup>R</sup> | This study |

### Sporulation and spore purification

Cells were grown in 25 ml liquid cell culture to OD_600_ 0.8 – 1.2. Glycerol was added to 1 M to induce sporulation. The liquid culture was incubated at 32 °C with vigorous shaking. After 24 h, spores and cells were harvested by centrifugation (10 min, 10,000 g and 25 °C). To further purify the wild-type spores, the pellet was resuspended in 5 ml water, the remaining vegetative cells were eliminated by sonication (Cole Palmer 4710 ultrasonic homogenizer, 30% output, 10 cycles), and sonication-resistant spores were washed three times with water and collected by centrifugation (5 min, 10,000 g and 4 °C). The elimination of vegetative cells was confirmed by DIC microscopy.

For phenotypic assays on starvation-induced fruiting body formation, vegetative cells (10 ul), at a concentration of 4 × 10^9^ ml^-1^, were spotted on CF (0.015% Casitone, 0.2% sodium citrate, 0.1% sodium pyruvate, 0.02% (NH_4_)_2_SO_4_, 10 mM MOPS (pH 7.6), 8 mM MgSO_4_ and 1 mM KH_2_PO_4_) plates containing an agar concentration of 1.5%, incubated at 32°C. To purify fruiting body spores, cells were grown in 25 ml liquid cell culture to OD_600_ 0.8 – 1.2 and harvested by centrifugation (10 min, 10,000 g and 25 °C). The pellet was resuspended in 1 ml water and plated on two 150 mm CF plates containing 1.5% agar using a spreader. The plates were air-dried and incubated at 32 °C for 120 h. For the wild-type and *ΔltgA* strains, fruiting bodies were scraped from the plates, suspended in 2 ml water, and subjected to sonication (Cole Palmer 4710 ultrasonic homogenizer, 60% output, 20 cycles). Spores were washed three times in water and collected by centrifugation (5 min, 10,000 g and 4 °C). For the *ΔltgB* strains, the sonication step was omitted.

### Quantification of starvation-induced spores

We used the submerge starvation culture to quantify starvation-induced spores. Cells were first grown in liquid CYE medium overnight and adjusted to OD_600_ 1 using fresh CYE. 50 ml of the asjusted culture was then poured to a Ø 150 mm polystyrene Petri dish and incubated without shaking for 24 h at 32 °C. Liquid and unattached cells were poured out carefully before adding 50 ml liquid CF medium. Petri dished were incubated without shaking for 96 h at 32 °C. Finally, liquid and unattached cells were poured out carefully and the cells attached to the bottom of Petri dished were scraped and resuspened into 1 ml water. The suspensions were then sonicated to eliminate vegetative cells, diluted and plated on solid CYE medium. Sonication-resistant spores were then numerated by counting the colonies after a 96-h incubation at 32 °C. Data were collected using three biological replicates.

### Peptidoglycan purification and UPLC analysis

For PG analysis, samples were processed as previously described for Gram negative bacteria (Alvarez *et al*., 2016, Desmarais *et al*., 2013). Vegetative cells were harvested at mid-stationary phase by centrifugation (30 min, 8,000 g). Vegetative cells and purified spores (as described in the previous section) were resuspended and boiled in 1x PBS with 5% SDS for 2 h. The sacculi were repeatedly washed with MilliQ water by ultracentrifugation (150,000 g, 10 min, 20 °C) to remove the remaining SDS. The samples were then treated with muramidase (100 µg/mL) for 16 hours at 37 °C. Muramidase digestion was stopped by boiling and coagulated proteins were removed by centrifugation (10 min, 15,000 g). The supernatants were first adjusted to pH 8.5-9.0 with sodium borate buffer and then sodium borohydride was added to a final concentration of 10 mg/ml. After reducing the samples at room temperature for 30 min, the pH was adjusted to pH 3.5 with orthophosphoric acid.

UPLC analyses of muropeptides were performed on a Waters UPLC system (Waters Corp., USA) equipped with an ACQUITY UPLC BEH C18 Column (Waters Corp., USA) and a dual wavelength absorbance detector. Elution of muropeptides was detected at 204 nm. Muropeptides were separated at 45 °C using a linear gradient from buffer A (0.1% formic acid in water) to buffer B (0.1% formic acid in acetonitrile) in a 25-min run, with a 0.30 ml/min flow.

### Muropeptide identification

Muropeptide identity was confirmed by MS and MS/MS analysis on a Xevo G2/XS Q-TOF mass spectrometer (Waters Corp., USA) coupled to the UPLC system with the same column, solvents, and gradients as mentioned above. Data acquisition and processing were performed using the UNIFI software package (Waters Corp., USA). Muropeptides were assigned based on: (i) accurate mass matching to theoretical monoisotopic masses of expected *M. xanthus* PG building blocks (Bui *et al*., 2009, White *et al*., 1968) and (ii) comparison of retention times with those reported in previous analysis with similar PG compositions. For vegetative cells analysis, relative total PG amounts were calculated by comparison of the total intensities of the chromatograms (total area) from three biological replicas normalized to the same initial biomass and extracted with the same volumes. Quantification of muropeptides was based on their relative abundances (relative area of the corresponding peak).

### Imaging and data analysis

For all imaging experiments on isolated spores/cells, we spotted 5 μl of cells grown in liquid CYE medium to OD_600_ ∼1 on agar (1.5%) pads and imaged using a Andor iXon Ultra 897 EMCCD camera (effective pixel size 160 nm) on an inverted Nikon Eclipse-Ti™ microscope with a 100✕ 1.49 NA TIRF objective. Fruiting bodies were photographed after a 120-h incubation on CF agar using a Nikon SMZ1000 microscope and an OMAX A3590U digital camera. Cell morphology was imaged at indicated time points using DIC microscopy. The geometric aspect ratios (L/W) of spores/cells were determined using a custom algorithm written in MATLAB, which is available in the GitHub repository, https://github.com/NanLabMyxo/Rod_shape_paper (Zhang *et al*., 2020, Zhang *et al*., 2023a, Zhang *et al*., 2023b).

For sptPALM, *M. xanthus* cells were grown in CYE to 4 ×10^8^ ml, spotted on 1.5% agar pads and subjected to highly inclined and laminated optical sheet (HILO) illumination (Fu *et al*., 2018, Nan *et al*., 2015, Nan *et al*., 2013, Tokunaga *et al*., 2008). PAmCherry was activated using a 405-nm laser (0.3 kW/cm^2^), excited and imaged using a 561-nm laser (0.2 kW/cm^2^). Images were acquired at 10 Hz. For each sptPALM experiment, single PAmCherry particles were localized in at least 100 individual cells from three biological replicates. sptPALM data were analyzed using a MATLAB (MathWorks) script , which is available in the GitHub repository, https://github.com/NanLabMyxo/Rod_shape_paper (Zhang *et al*., 2020, Zhang *et al*., 2023a, Zhang *et al*., 2023b, Ramírez Carbó & Nan, 2026). Briefly, cells were identified using differential interference contrast images. Single PAmCherry particles inside cells were fit by a symmetric 2D Gaussian function, whose center was assumed to be the particle’s position (Fu *et al*., 2018). Particles that explored areas smaller than 160 nm × 160 nm (within one pixel) in 0.4 - 1.2 s were considered immotile (Fu *et al*., 2018, Zhang *et al*., 2023a, Zhang *et al*., 2023b). *D* of all the mobile particles was determined from a linear fit to the first four points of the MSD using a formula MSD = 4*DΔt* (Fu *et al*., 2018, Lee *et al*., 2016). Error bars were the standard derivation of 1,000 bootstrap samples using the published method (Morgenstein *et al*., 2015). Sample trajectories were generated using the TrackMate (Ershov *et al*., 2022) plugin in the ImageJ suite (https://imagej.net).

### Immunofluorescence

Cell samples were resuspended and boiled in 1x PBS with 5% SDS for 2 h. The sacculi were repeatedly washed with water by centrifugation (5 min, 15,000 g and 25 °C) to remove the remaining SDS. Remaining proteins were further removed by incubation with proteinase K (1 µg/ml) for 2 h at 37 °C. Microscope cover slides were prepared by coating with 100 µl 0.1% poly-L-lysine for 5 min at room temperature (RT), washing with ddH_2_O and air drying. 100 µl samples were spotted onto the slides and incubated for 20 min at RT. The slides were then washed three times with phosphate buffer saline (PBS; 81 mM Na_2_HPO_4_, 15 mM KH_2_PO_4_, 1.37 M NaCl, 27 mM KCl, pH 7.4), and then incubated with anti-PG primary antibody^16^ diluted 1:1000 in PBS at RT for 1 h. Slides were washed 6 times with PBS and then incubated with Alexa Fluor 488 conjugated goat-anti-rabbit secondary antibody (Invitrogen) at a 1:1000 dilution in PBS at RT for 1 h in the dark. Slides were washed 6 times with PBS. Each cover glass was applied to a supporting slide, fixed with nail polish, and stored at 4 °C, if necessary. Images were recorded using a 488-nm laser.

### Protein expression and purification

DNA sequences encoding amino acids 21 – 243 of LtgA and 26 - 710 of LtgB were amplified by polymerase chain reaction (PCR) and inserted into the pET28a vector (Novagen) between the restriction sites of *Eco*RI and *Hind*III and used to transform *E. coli* strain BL21(DE3). Transformed cells were cultured in 20 ml LB (Luria-Bertani) broth at 37 °C overnight and used to inoculate 1 L LB medium supplemented with 1.0% glucose. Protein expression was induced by 0.1 mM IPTG (isopropyl-h-d-thiogalactopyranoside) when the culture reached an OD_600_ of 0.8. Cultivation was continued at 16 °C for 10 h before the cells were harvested by centrifugation at 6,000 × g for 20 min and lysed by sonication in buffer A (20 mM Tris-HCl pH 8.0, 200 mM NaCl), (Nan *et al*., 2010, Nan *et al*., 2006). Proteins were loaded to an NGC™ Chromatography System (BIO-RAD) and 5-ml HisTrap™ columns (Cytiva) and eluted by buffer B (20 mM Tris-HCl pH 8.0, 200 mM NaCl, 500 mM immidazole) (Pogue *et al*., 2018, Nan *et al*., 2010). Purified proteins were concentrated using Amicon™ Ultra centrifugal filter units (Millipore Sigma) with a 10-kDa molecular weight cutoff and stored at -80 °C.

### LTG activity (RBB) assay

PG was purified following the published method (Zhang *et al*., 2023b, Alvarez *et al*., 2016, Ramirez Carbo *et al*., 2024). In brief, *M. xanthus* cells were grown until mid-stationary phase and harvested by centrifugation (6,000 × g, 20 min, 25 °C). Supernatant was discarded and the pellet was resuspended and boiled in 1× PBS with 5% SDS for 2 h. SDS was removed by repetitive wash with water and centrifugation (21,000 × g, 10 min, 25 °C). Purified PG from 100 ml culture was suspended into 1 ml 1× PBS and stored at -20 °C. RBB labelling of PG was performed essentially as previously described (Uehara *et al*., 2010, Jorgenson *et al*., 2014). Purified sacculi were incubated with 20 mM RBB in 0.25 M NaOH overnight at 37 °C. Reactions were neutralized by adding equal volumes of 0.25 M HCl and RBB-labeled PG was collected by centrifugation at 21,000 × g for 15 min. Pellets were washed repeatedly with water until the supernatants became colorless. RBB-labelled sacculi were incubated with purified LtgA or LtgB (1 mg/ml) at 25 °C for 12 h. lysozyme (1 mg/ml) was used as a positive control. Dye release was quantified by the absorption at 595 nm from the supernatants after centrifugation (21,000 × g, 10 min, 25 °C).

### Immunoblotting

Cells were grown in liquid CYE medium overnight, harvested, and concentrated to OD_600_ 10. 10 µl of cell suspension was mixed with 10 µl of loading buffer and 10 µl of the mixture was subjected to SDS-PAGE. The expression and stability of PAmCherry-labeled proteins were determined by immunoblotting using an anti-mCherry antibody (Rockland Immunochemicals, Inc., Lot 46705) and a goat anti-Rabbit IgG (H+L) secondary antibody, HRP (Thermo Fisher Scientific, catalog # 31460). MreB was detected as the loading control using an anti-MreB serum(Mauriello *et al*., 2010) and the same secondary antibody. The blots were developed with Pierce™ ECL Western Blotting Substrate (Thermo Fisher Scientific REF 32109) and a MINI-MED 90 processor (AFP Manufacturing).

## Supporting information

Sporulation manuscript V3

## Acknowledgements

This work was supported by the National Institutes of Health grants GM129000 to B. N. Research in the F. C. laboratory is supported by the Swedish Research Council, the Laboratory for Molecular Infection Medicine Sweden (MIMS), Umeå University, the Knut and Alice Wallenberg Foundation (KAW) and the Kempe Foundation. B. N.’s group received financial support from Dr. David R. Zusman, who played no role in the design, execution, or presentation of this work.

