## Supplementary material for "A novel mechanism for morphological change during bacterial sporulation based on programmed peptidoglycan degradation": Sporulation manuscript V3

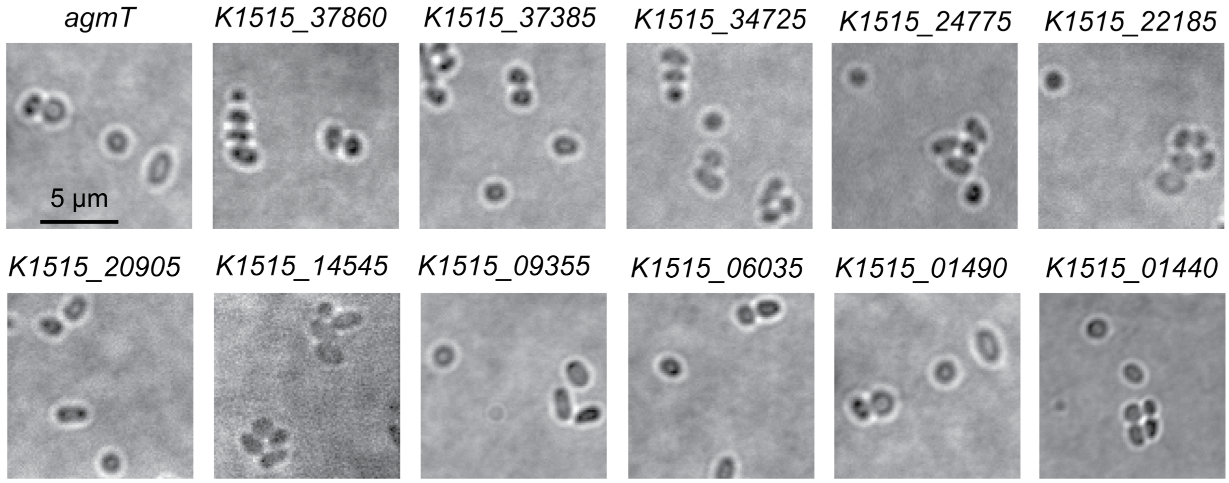


**Figure 2 – figure supplement 1. Besides LtgA and LtgB, other putative LTGs do not affect either sporulation pathway significantly.** Bright field images of cells were obtained after 4 h of glycerol-induction.

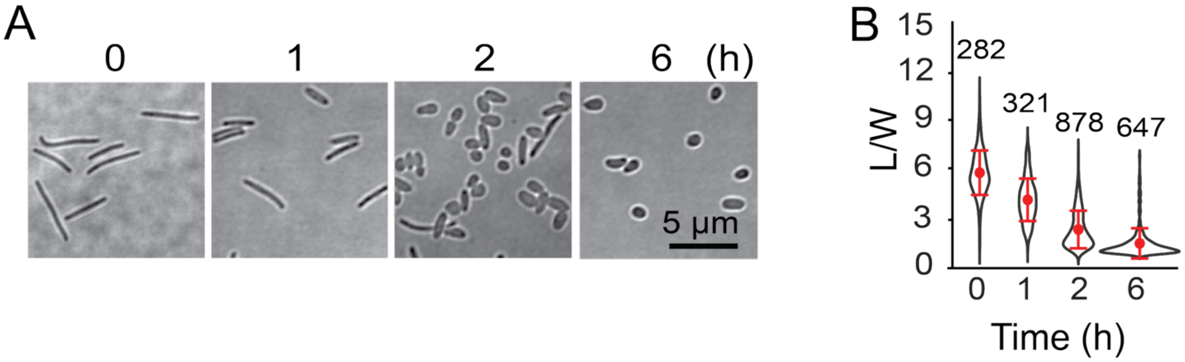


**Figure 2 – figure supplement 2.** In the absence of vanillate, $P_{van}$-driven MurA expression does not affect glycerol-induced sporulation. **A)** Bright field images of cells at different time points after glycerol-induction. **B)** Quantitative analysis of glycerol-induced sporulation using the length/width ratio (L/W) of cells. Whiskers indicate the 25th - 75th percentiles and red dots the median. The total number of cells analyzed is shown on top of each plot.

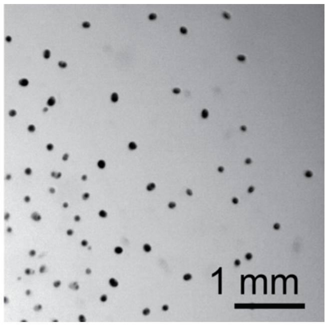


**Figure 3 – Figure supplement 1.** In the absence of vanillate, $P_{van}$-driven MurA expression does not affect starvation-induced fruiting body formation. The image was taken after 96 h of incubation on a CF agar.

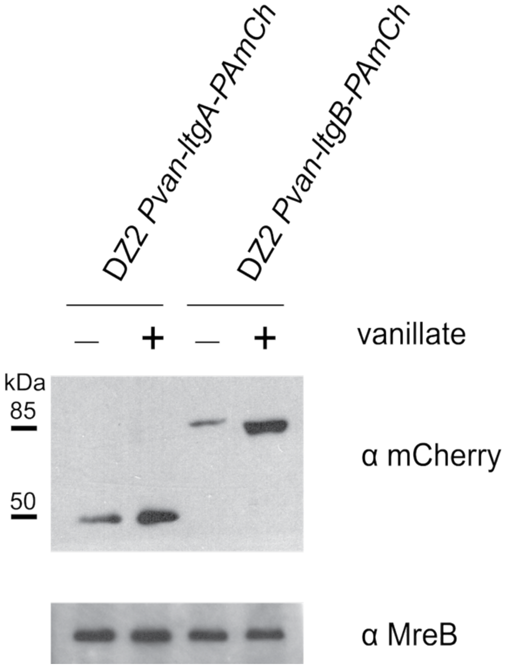


**Figure 4 – figure supplement 1. Vanillate-induced expression of LtgA-PAmCherry and LtgB-PAmCherry.** Protein expression was visualized through immunoblotting using mCherry antibodies (α mCherry). The MreB protein was used as the loading control, detected by the MreB antibodies (α MreB). "—", no vanillate; "+", 200 μM sodium vanillate.

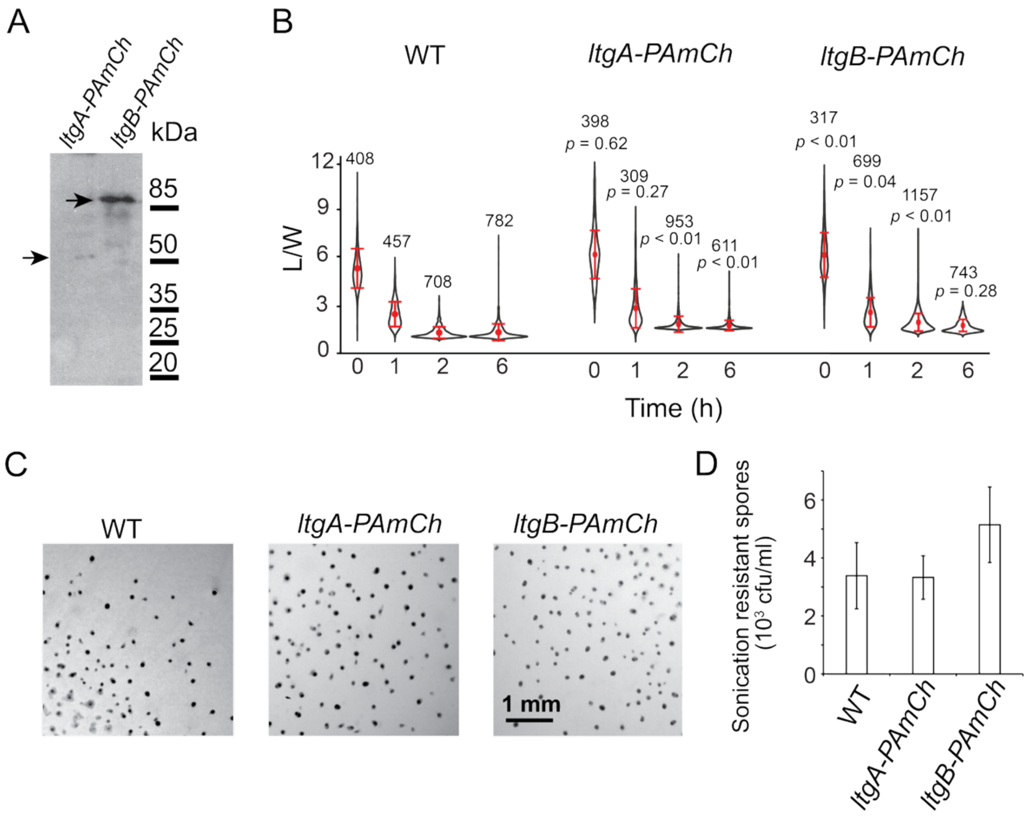


**Figure 5 – figure supplement 1. PAmCherry-labeled LtgA and LtgB are fully functional. A)** PAmCherry-labeled LtgA and LtgB are expressed as full-length proteins in vegetative cells (53 kDa and 103 kDa, respectively), which were detected by an anti-mCherry antibody. **B)** The PAmCherry labels do not affect glycerol-induced sporulation. Quantitative analysis of glycerol-induced sporulation using the length/width ratio (L/W) of cells. Whiskers indicate the 25th - 75th percentiles and red dots the median. The total number of cells analyzed is shown on top of each plot. **C**) The PAmCherry labels do not inhibit starvation-induced fruiting body formation on agar surfaces. The images were taken after 96 h of incubation on CF agar. **D)** The PAmCherry labels do not inhibit the formation of starvation-induced spores. Sonication-resistant spores were counted as colony formation units (cfu) after 96 h of incubation on CYE agar. Data are shown as mean ± standard deviation (n = 3).
